# Mitotic replisome disassembly in vertebrates

**DOI:** 10.1101/418368

**Authors:** Sara Priego Moreno, Rebecca M. Jones, Divyasree Poovathumkadavil, Agnieszka Gambus

## Abstract

Recent years have brought a breakthrough in our understanding of the process of eukaryotic DNA replication termination. We have shown that the process of replication machinery (replisome) disassembly at the termination of DNA replication forks in S-phase of the cell cycle is driven through polyubiquitylation of one of the replicative helicase subunits Mcm7. Our previous work in *C.elegans* embryos suggested also an existence of a back-up pathway of replisome disassembly in mitosis. Here we show, that in *Xenopus laevis* egg extract, any replisome retained on chromatin after S-phase is indeed removed from chromatin in mitosis. This mitotic disassembly pathway depends on formation of K6 and K63 ubiquitin chains on Mcm7 by TRAIP ubiquitin ligase and activity of p97/VCP protein segregase. The mitotic replisome pathway is therefore conserved through evolution in higher eukaryotes. However, unlike in lower eukaryotes it does not require SUMO modifications. This process can also remove any helicases from chromatin, including “active” stalled ones, indicating a much wider application of this pathway than just a “back-up” for terminated helicases.

## INTRODUCTION

Faithful cell division is the basis for the propagation of life and requires accurate duplication of all genetic information. DNA replication must be precisely regulated as unrepaired mistakes can change cell behaviour with potentially severe consequences, such as genetic disease, cancer and premature ageing (Burrell, McClelland et al., 2013). Fundamental studies have led to a step-change in our understanding of the initiation of DNA replication and DNA synthesis, but until discovery of the first elements of the eukaryotic replisome disassembly mechanism in 2014, the termination stage of eukaryotic replication was mostly unexplored.

DNA replication initiates from thousands of replication origins. They are the positions within the genome where replicative helicases become activated and start unwinding DNA while moving in opposite directions, away from each other, creating two DNA replication forks. The replicative helicase is composed of Cdc45, Mcm2-7 hexamer and GINS complex (CMG complex) (Moyer, Lewis et al., 2006); it is positioned at the tip of replication forks and forms a platform for replisome assembly (Replisome Progression Complex, RPC) (Gambus, Jones et al., 2006). Once established, the replication forks replicate chromatin until they encounter forks coming in opposite directions from neighboring origins. At this point the termination of replication forks takes place. As CMG helicases travel on the leading strand templates at the forks, the strand encircled by converging helicases differs due to the antiparallel nature of the DNA molecule (Fu, Yardimci et al., 2011). The two converging helicases can therefore pass each other allowing for completion of DNA synthesis. Finally, removal of the replisome from fully duplicated DNA is the last stage of forks termination (Dewar, Budzowska et al., 2015). We have shown that in *Xenopus laevis* egg extract and in *C. elegans* embryos this replisome removal in S-phase is driven by Cul2^LRR1^ ubiquitin ligase, which ubiquitylates Mcm7 within the terminated CMG complex (Sonneville, Moreno et al., 2017). Such modified CMG is then recognized by p97 (VCP) segregase and removed from chromatin allowing for disassembly of the whole replisome built around the helicase (Moreno, Bailey et al., 2014).

Interestingly, we have shown that in *C. elegans* embryos, any helicase complexes that fail to be unloaded in S-phase are alternatively unloaded in prophase of mitosis (Sonneville et al., 2017). This potential back-up mechanism functions when CUL-2^LRR-1^ activity is blocked and, like the S-phase pathway, depends on the p97 segregase for unloading. Unlike the S-phase pathway, however, it requires an additional p97 cofactor UBXN-3/FAF1 and the SUMO-protease ULP-4 (Senp6/7 homologue in higher eukaryotes) (Sonneville et al., 2017). Interestingly, budding yeast do not possess this mitotic replisome disassembly pathway; cells lacking SCF^Dia2^ activity, the ubiquitin ligase responsible for Mcm7 ubiquitylation in *S. cerevisiae*, accumulate post-termination replisomes on DNA until the next G1 of the next cell cycle (Maric, Maculins et al., 2014). Our aim therefore was to determine if this mitotic replisome disassembly pathway has been conserved throughout evolution and is functioning in higher eukaryotes, or if it is a phenomenon specific to *C. elegans* embryos. Here we show that a mitotic disassembly pathway does exist in vertebrates and determine the first elements of its regulation. We show that only a restricted part of the replisome stays retained on chromatin throughout mitosis in *Xenopus* egg extract. The disassembly of this replisome is independent of cullin type ubiquitin ligases but requires p97 segregase function. Mitotic replisome disassembly depends on K6 and K63 ubiquitin chains but not SUMO modifications. We also identify TRAIP ubiquitin ligase as essential for Mcm7 ubiquitylation in mitosis. Finally, we show that active forms of helicase can also be unloaded in this way, suggesting that rather than being a back-up pathway for the disassembly of terminated replisomes, this process is essential to remove any replisome from chromatin before cell division.

## RESULTS

*Xenopus* egg extract is a cell-free system, which has proven to be instrumental over the years in studies of DNA replication. To determine the existence of a mitotic replisome disassembly pathway in *Xenopus* egg extract we needed to allow our extract to progress from interphase into mitosis. Routinely we block this cell cycle transition by inhibition of cyclin synthesis in the extract through supplementation with cycloheximide, since it allows for better synchronization of the replication reaction (Gillespie, Gambus et al., 2012). To allow the extract to progress into mitosis we need therefore to supplement it with cyclins – either by addition of mitotic extract or addition of recombinant cyclin. We purified His-tagged *Xenopus laevis* Cyclin A1 NΔ56 (hereafter: Cyclin A1Δ) and added it to the extract upon completion of DNA replication, as described previously, to induce mitotic entry (EV Fig 1A) (Strausfeld, Howell et al., 1996). Upon addition of Cyclin A1Δ we could see condensation of chromatin into chromosomes – a clear sign of mitosis (EV Fig 1B).

To test if the replisome retained on chromatin in S-phase can be unloaded as cells enter mitosis we followed the experimental path detailed in Fig 1A. A replication reaction was set up in the interphase extract supplemented with Cullin ligase inhibitor MLN4924 to block Cul2^LRR1^ activity and the S-phase replisome unloading (Sonneville et al., 2017). After completion of replication (90 min, EV Fig 1A) we optionally added Cyclin A1Δ, isolated chromatin at different time points during mitosis progression and analysed chromatin bound proteins by western blotting (Fig 1B). In the late S-phase samples (buffer control), the CMG helicase (represented by Cdc45 subunit) remained associated with chromatin and Mcm7 displayed low levels of ubiquitylation, as seen before upon MLN4924 treatment and Cul2 immunodepletion (Moreno et al., 2014, Sonneville et al., 2017), however upon addition of Cyclin A1Δ, Cdc45 and ubiquitylated Mcm7 were efficiently unloaded. This result indicates that indeed the mitotic replisome disassembly pathway is evolutionarily conserved and exists in vertebrates and that, unlike the S-phase pathway, it does not require activity of Cullin type ubiquitin ligases because the Cullin inhibitor was present throughout the reaction (Fig 1B).

**Figure 1.**
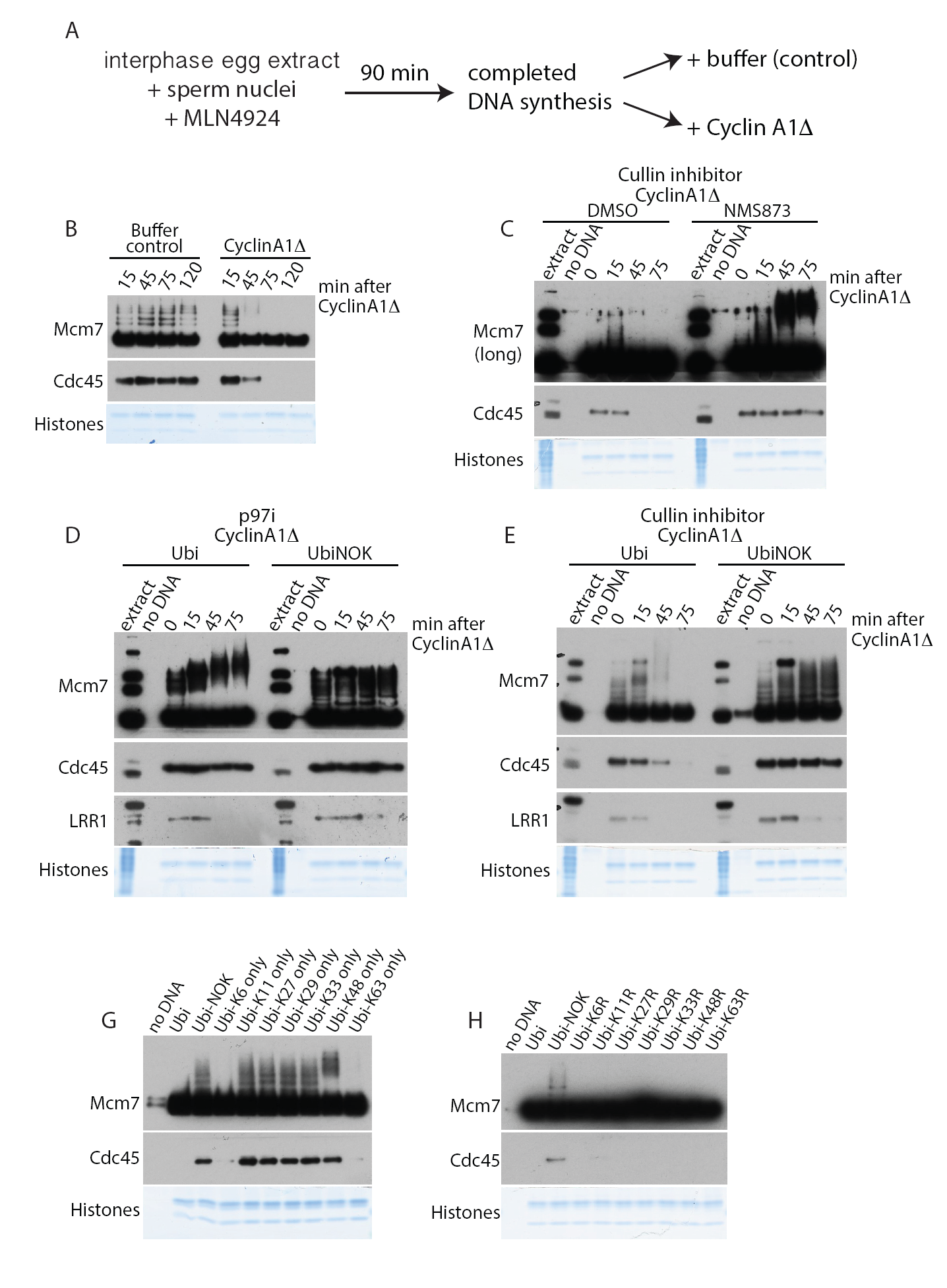
Mcm7 is ubiquitylated with K6 and K63 ubiquitin chains in mitosis and removed from chromatin by p97 segregase. **(A)** Experimental design for driving egg extract into mitosis. **(B)** Experiment following design in (A). DNA was replicated to completion in egg extract supplemented with Cullin inhibitor MLN4924. After 90 min of replication reaction Cyclin A1Δ was optionally added to the extract to drive extract to mitosis. Chromatin was isolated at indicated time-points after Cyclin A1Δ addition and chromatin samples analysed by western blotting with indicated antibodies. Colloidal Coomassie stained histones serve as a quality and loading control. **(C)** The replication reaction was completed in the presence of Cullin inhibitor MLN4924 and driven into mitosis by addition of Cyclin A1Δ. At the same time as Cyclin A1Δ, half of the sample was supplemented additionally with p97 inhibitor NMS873. Chromatin samples were isolated at indicated time-points and analysed as in (B). A sample without DNA addition (no DNA) was processed alongside others as a chromatin specificity control. **(D)** The replication reaction was completed in the presence of p97 inhibitor NMS873 and driven into mitosis by addition of Cyclin A1Δ. At the same time as Cyclin A1Δ, samples were supplemented with recombinant wt ubiquitin or UbiNOK. Chromatin samples were analysed as above. **(E)** Experiment as in (D) but replication reaction was carried out in the presence of Cullin inhibitor MLN4924 instead of p97 inhibitor. **(G)** and **(H)** Replication reaction was completed in the presence of Cullin inhibitor MLN4924 and driven to mitosis by addition of Cyclin A1Δ. At the same time as Cyclin A1Δ addition, extract was supplemented with indicated mutants of ubiquitin. Chromatin was isolated at 75 minutes after Cyclin A1Δ addition and analysed by western blotting as above.

Next, we tested whether the mitotic replisome disassembly pathway required the activity of the p97 segregase. We followed the experimental setup as before but now optionally added the inhibitor of p97, NMS873. The retained replisome was unloaded upon Cyclin A1Δ addition in the absence but not in the presence of the p97 inhibitor indicating that indeed p97 does play an essential role in promoting mitotic replisome disassembly (Fig 1C). We could also see an analogous result if the p97 inhibitor was present throughout the two stages of the cell cycle as the only way to block replisome disassembly (EV Fig 1C).

Interestingly, when the mitotic unloading of replisome was blocked with p97 inhibitor, we could clearly see accumulation of highly modified forms of Mcm7 on chromatin (Fig 1C and EV Fig 1C). To examine whether these modifications were due to further ubiquitylation of Mcm7, we blocked S-phase and mitotic replisome disassembly by addition of p97 inhibitor from the beginning of the replication reaction, induced mitosis after completion of DNA sythesis and optionally supplemented extract with a high concentration of wild-type (wt) ubiquitin or a chain-terminating mutant of ubiquitin with all lysines mutated (UbiNOK).

Supplementation of mitotic extract with wt ubiquitin allowed for accumulation of highly modified Mcm7 on chromatin in mitosis as before (Fig 1D, left). However, addition of UbiNOK blocked further modifications of Mcm7, leaving only the chains which were built previously in S-phase (Fig1D, right). To determine whether this further Mcm7 polyubiquitylation in mitosis was essential for mitotic replisome disassembly we repeated the experiment with addition of wt Ubi or UbiNOK but this time only in the presence of the Cullin inhibitor from the start of the reaction (Fig 1E). Addition of UbiNOK to mitotic extract blocked disassembly of the replisome (as shown by permanent Cdc45 chromatin binding) suggesting that further Mcm7 polyubiquitylation is required for its unloading. We could also observe that LRR1 (the substrate specific subunit of Cullin 2, targeting Mcm7 in S-phase) dissociates from chromatin in mitosis irrespectively of replisome disassembly, in agreement with the finding that it does not play an essential role in this pathway (Fig 1D and E).

As the ubiquitin ligase acting in the mitotic pathway differed from that of the S-phase pathway, we decided to test whether the type of ubiquitin chains built on Mcm7 in mitosis also differed. To determine which ubiquitin chains were required for the mitotic Mcm7 ubiquitylation and replisome disassembly, we supplemented extract with Cullin inhibitor, allowed completion of DNA synthesis and subsequently induced mitosis along with addition of a series of ubiquitin mutants that have only one lysine left in their sequence (Fig 1G). We observed that only wt ubiquitin and ubiquitin containing lysine 6 (K6) or lysine 63 (K63) could support mitotic replisome disassembly (as visualised by absence of Cdc45 on chromatin at 75 min after inducing mitosis) (Fig 1G). Interestingly, chains linked through lysine 48 (K48), which are responsible for S-phase unloading (Moreno et al., 2014), could still be attached to Mcm7 in mitosis (upshift of modified Mcm7 forms) but they could not support unloading of the replisome as Cdc45 remained associated with chromatin. In a reciprocal experiment, we used a series of ubiquitin mutants with only one of the lysines within ubiquitin mutated (Fig 1H). All of the mutants used, apart from the UbiNOK control mutant, supported disassembly of the replisome, suggesting that either K6 or K63 can fulfill the mitotic pathway requirements (Fig 1H).

Having established that the type of ubiquitin chains and the type of ubiquitin ligase used by the mitotic pathway were different to those used by the S-phase pathway, our aim was to determine which ligase is driving this mitotic disassembly pathway. To identify the potential ligase we decided to immunoprecipitate the replisome retained on mitotic chromatin and analyse all the interacting proteins by mass spectrometry. We set up a replication reaction in the presence of caffeine and the p97 inhibitor and induced mitosis upon completion of DNA synthesis. We immunoprecipitated Mcm3 from mitotic chromatin and analysed interacting factors by mass spectrometry. Firstly, we determined which components of the replisome are still retained on chromatin in mitosis. For this we compared the replisome components retained on chromatin in mitosis with the S-phase post-termination replisome, reported previously (Sonneville et al., 2017) (Fig 2A and B). Interestingly, while inhibition of replisome disassembly in S-phase led to accumulation of the whole replisome on chromatin (Sonneville et al., 2017), only a selection of replisome components stayed on chromatin in mitosis. All of the lagging strand components of the replisome were lost, as were Mcm10 and Claspin, while levels of Ctf4/And-1, Timeless/Tipin and Pol epsilon were also reduced (Fig 2A and B). This suggests that only components directly interacting with the CMG accumulated around it through to mitosis, while others, more peripheral to CMG could dissociate over time.

**Figure 2.**
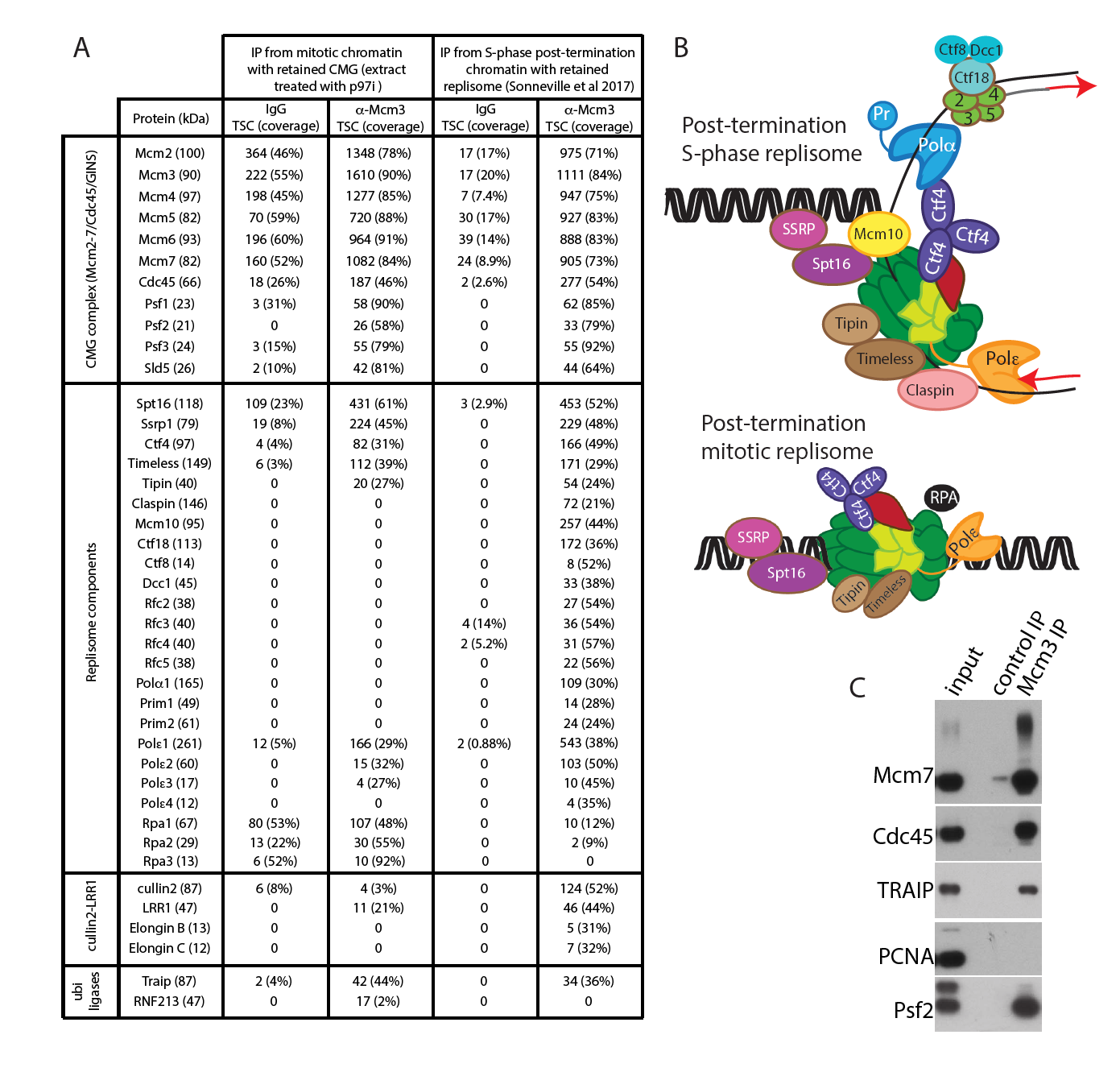
Composition of the replisome retained on mitotic chromatin. **(A)** Replication reaction was completed in egg extract supplemented with caffeine and p97 inhibitor NMS873, extract was driven into mitosis by addition of Cyclin A1Δ, chromatin isolated at 60 minutes after Cyclin A1Δ addition and chromatin proteins released from DNA. Antibodies against Mcm3 (or control IgG) were used to immunoprecipitate replisomes and immunoprecipitated samples were analysed by mass spectrometry. The total spectral count for each identified replisome component is presented together with sequence coverage of analysed peptides. The result of analysis of mitotic retained replisome is compared with S-phase post-replication replisome reported in (Sonneville et al., 2017). **(B)** Schematic representation of the data presented in (A). **(C)** A small proportion of the material from the mitotic Mcm3 IP experiment in (A) was analysed by western blotting with indicated antibodies.

The level of histone chaperone FACT (Spt16 and SSRP) stays the same between S-phase and mitosis. This suggests that the retained replisome has the potential ability to move through chromatin as FACT is likely to displace nucleosomes in front of such a replisome. We could see also that Cul2^LRR1^, which strongly accumulated in the S-phase post-termination replisome, is not a major component of the mitotic replisome, as expected from previous data (Fig 1).

Finally, we detected two other ubiquitin ligases interacting with the mitotic helicase: TRAIP and RNF213. More specifically, we found that TRAIP interacts with the post-termination replisome in S-phase but it is enriched in mitosis, while RNF213 is a minor interactor of only the mitotic replisome (Fig 2A). The TNF-receptor-associated factor (TRAF)-interacting protein (TRAIP, also known as TRIP or RNF206) was originally identified through its ability to bind TRAF1 and TRAF2 and shown to inhibit NFkB activation (Lee, Lee et al., 1997). It has been since shown that TRAIP is an E3 ubiquitin ligase, which is essential for cell proliferation (Besse, Campos et al., 2007, Park, Choi et al., 2007), and which is required for resolution of replication stress (Feng, Guo et al., 2016, Harley, Murina et al., 2016, Hoffmann, Smedegaard et al., 2016) and for regulation of the spindle assembly checkpoint during mitosis (Chapard, Meraldi et al., 2014). TRAIP is ubiquitously expressed, with its expression regulated by E2F transcription factors and protein stability controlled by the ubiquitin proteasome pathway – as a result, the protein level of TRAIP peaks in the G2/M stage of the cell cycle (Chapard, Hohl et al., 2015). On the other hand, RNF213 (mysterin) is a large (591kD) ATPase/E3 ligase, which is mostly known as being a susceptibility gene for Moyamoya disease (MMD) (cerebrovascular disease) (Kamada, Aoki et al., 2011, Liu, Morito et al., 2011). Of note, *RNF213-/*-mice do not show any apparent health problems (Kobayashi, Yamazaki et al., 2013, Sonobe, Fujimura et al., 2014) and more recently RNF213 was shown to globally regulate (α-ketoglutarate)-dependent dioxygenases (α-KGDDs) and non-mitochondrial oxygen consumption (NMOC) (Banh, Iorio et al., 2016). To support our mass spectrometry data, we have tested a number of antibodies by western blotting against RNF213 and TRAIP to confirm their association with the chromatin bound replisome in mitosis. Whilst we were unsuccessful with detection of any signal for RNF213, we were able to show that TRAIP interacts with replisome retained on chromatin in mitosis (Fig 2C and EV Fig1D).

As we confirmed that TRAIP is a likely candidate for the ubiquitin ligase ubiquitylating Mcm7 and leading to disassembly of the replisome retained in mitosis, we next characterized TRAIP chromatin binding dynamics during the two cell cycle stages and the replisome disassembly process. TRAIP associated weakly with S-phase chromatin at times when forks are moving through chromatin replicating DNA. However, it accumulated strongly on chromatin in S-phase upon inhibition of replisome disassembly with the p97 inhibitor (Fig 3A). Importantly, TRAIP also accumulated on mitotic chromatin when replisome disassembly was inhibited with the p97 inhibitor, following the same pattern as replisome components (Fig 3 B). To test whether TRAIP is the ubiquitin ligase ubiquitylating Mcm7 in mitosis, we purified recombinant GST-tagged *X.laevis* TRAIP, both wild type and C25A RING domain mutant, which been shown to disrupt TRAIP ubiquitin ligase activity (Besse et al., 2007, Chapard et al., 2014). We blocked the disassembly of the replisome in S-phase by inhibition of cullins with MLN4924, and drove extract into mitosis by addition of Cyclin A1Δ, when we added recombinant wt or mutant TRAIP. To be able to clearly see ubiquitylation of Mcm7 in mitosis we supplemented the mitotic extract also with p97 inhibitor NMS873 to inhibit unloading of ubiquitylated Mcm7. Figure 3C shows that addition of RING mutant of TRAIP, but not wild type TRAIP, inhibits mitotic ubiquitylation of Mcm7 as the size of ubiquitylated forms of Mcm7 remains very close to ones build on Mcm7 already in S-phase (15 min timepoint). This suggest that the recombinant mutant TRAIP successfully competed with endogenous TRAIP protein and that ubiquitin ligase activity of TRAIP is needed for mitotic Mcm7 ubiquitylation.

**Figure 3.**
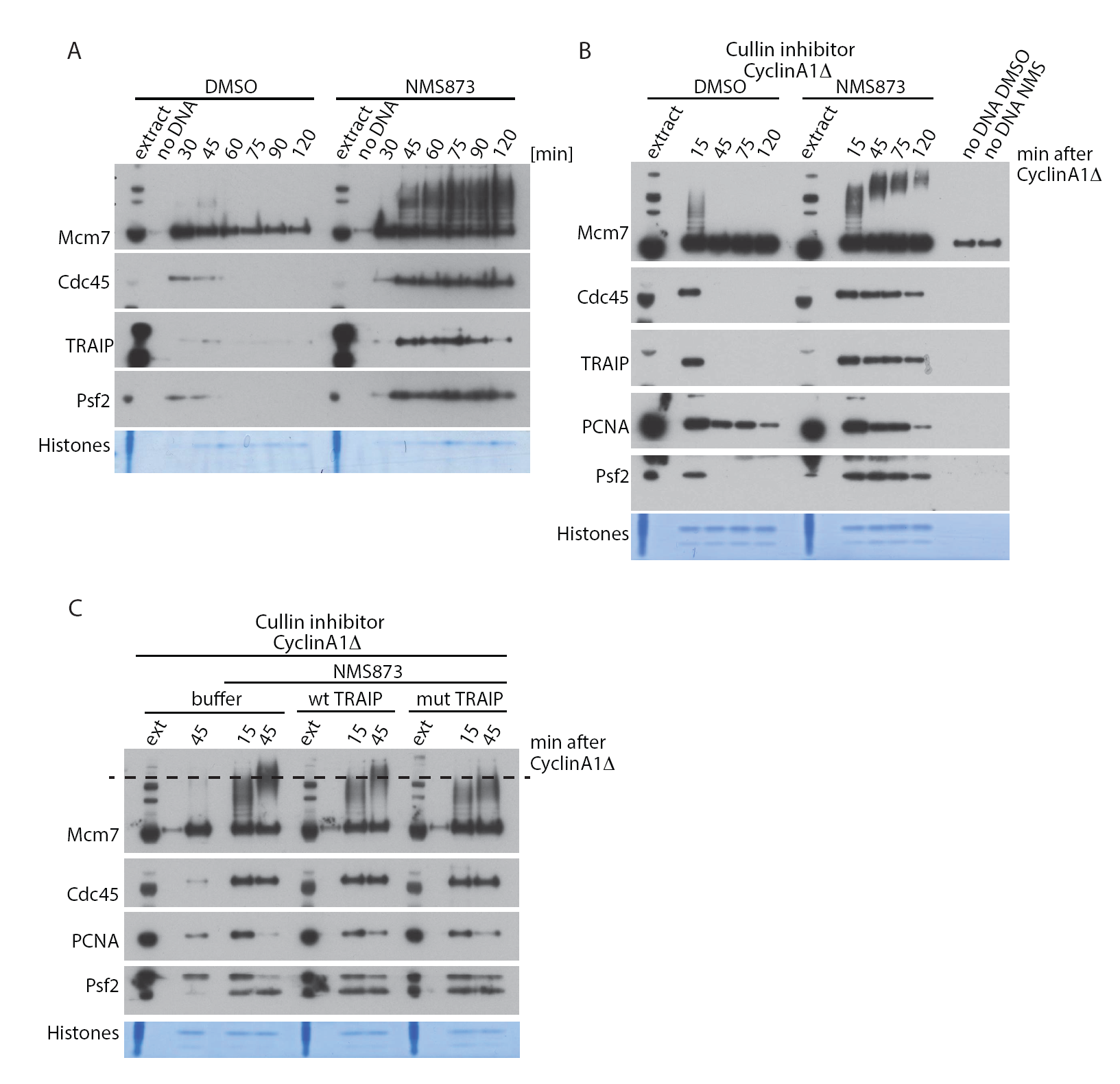
TRAIP regulates replisome disassembly in mitosis. **(A)** Sperm DNA was replicated in egg extract optionally supplemented with p97 inhibitor NMS873. Chromatin samples were isolated during reaction at indicated time-points and analysed as in Fig 1. **(B)** Experiment analogous to Fig 1C but analysed with indicated antibodies. **(C)** The replication reaction was completed in the presence of Cullin inhibitor MLN4924 and driven into mitosis by addition of Cyclin A1Δ. At the same time as Cyclin A1Δ, most of the samples were supplemented with p97 inhibitor NMS873 and optionally with LFB1/50 buffer or wt TRAIP or RING mutant (C25A) TRAIP at 30 μg/ml. Chromatin samples were isolated at indicated time-points and analysed with indicated antibodies. The dashed line on Mcm7 blot runs through the middle of ubiquitylation signal for Mcm7 in mitosis in control (buffer) sample to aid comparison between samples.

To fully understand the requirement for ubiquitin like modifications during the mitotic replisome disassembly in vertebrates, we aimed to establish whether SUMOylation plays any role in this process as ULP-4 is essential for mitotic helicase disassembly in *C. elegans* embryos. To this end we decided to inhibit or stimulate SUMOylation during mitosis and assess its effect on replisome disassembly. Firstly, we observed that late S-phase chromatin is full of SUMOylated factors and that levels of these proteins go down over time upon entry into mitosis (Fig 4 and EV Fig 2). To inhibit SUMOylation we have supplemented the mitotic extract with the recombinant active domain of SENP1 which acts as a potent non-specific deSUMOylating enzyme. Addition of SENP1 wipes out all the SUMO2ylation (Fig 4A) and SUMO1ylation (EV Fig 3A) but the disassembly of the mitotic replisome is not affected (Fig 4A). We have also stimulated SUMOylation through addition of a high concentration of recombinant SUMO1 or SUMO2 (EV Fig 2). In both cases, despite a clear increase of SUMO signal on chromatin, the unloading of the mitotic replisome was not affected. Finally we have also blocked de-SUMOylation with SUMO2-VS, a derivative of SUMO2 which binds to the active site of SENPs and blocks their activity. Again, we could observe strong accumulation of SUMO2ylated products in the extract and on chromatin without affecting mitotic replisome disassembly. Interestingly, despite inhibition of de-SUMOylating enzymes, most of the SUMO signal is still disappearing from chromatin during progression of mitosis, indicating that the SUMOylated proteins are unloaded from chromatin throughout mitosis rather than being de-SUMOylated. In conclusion, we determined that SUMO modifications do not play an essential role in the mitotic disassembly pathway in *Xenopus* egg extract. In an analogous way we have also shown that they do not play a role during the S-phase replisome disassembly pathway (EV Fig 3).

**Figure 4.**
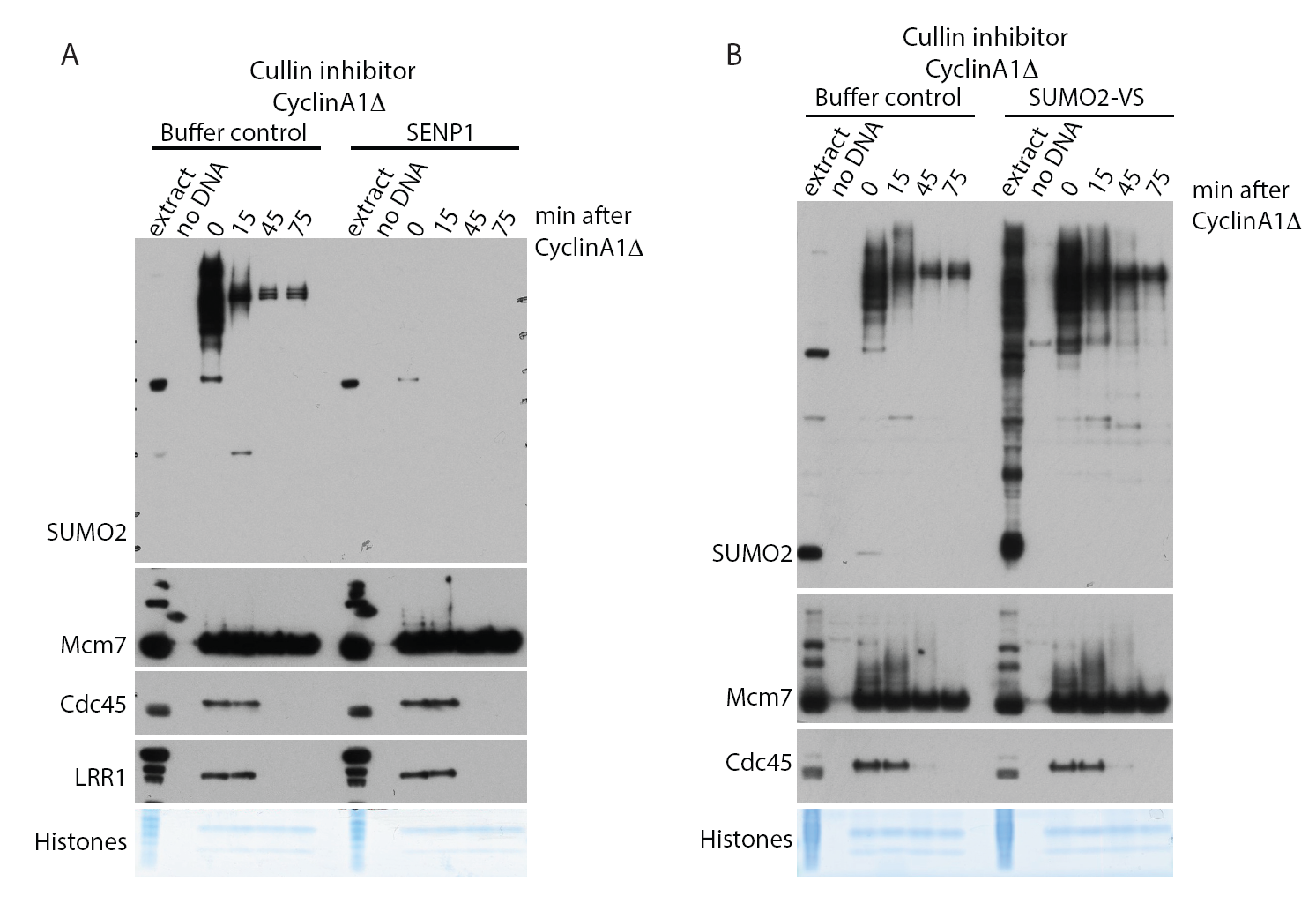
SUMOylation is not required for mitotic replisome disassembly. **(A)** The replication reaction was completed in the presence of Cullin inhibitor MLN4924 and driven into mitosis by addition of Cyclin A1Δ. At the same time as Cyclin A1Δ, half of the sample was supplemented additionally with active domain of SENP1. Chromatin samples were isolated at indicated time-points and analysed as in Fig 1. **(B)** As in (A) but instead of supplementing extract with SENP1 we supplemented it with SENPs inhibitor SUMO2-VS.

Finally, we set out to determine whether the mitotic replisome disassembly pathway we were characterising was a mere “back-up” pathway for replisomes that terminated in S-phase but failed to be unloaded, or if it can have a more generic usage to remove any replication machinery still remaining on chromatin until mitosis. To test such a possibility, we blocked active (stalled) replisomes on chromatin by addition of polymerase inhibitor aphidicolin to the egg extract during DNA replication reaction. To accumulate such replisomes in large numbers, we also supplemented the extract with caffeine so as to block checkpoint activation and fire origins uncontrollably. Upon accumulation of such blocked replisomes we supplemented the reaction optionally with Cyclin A1Δ at 90 min to induce mitotic entry (Fig 5A). Interestingly, active replisomes remained associated with chromatin throughout the experiment in late S-phase (buffer control), with no indication of Mcm7 ubiquitylation as expected. Upon addition of cyclin A1Δ however, Mcm7 becomes ubiquitylated and replisomes are unloaded (Cyclin A1Δ). We can observe however, a noticeable delay of both these processes compared with terminated replisomes (compare Fig 5A with Fig 1B). Such a delay is likely due to the fact that with no prior ubiquitylation of Mcm7 in S-phase, it takes longer for ubiquitin chains to be built in mitosis. We have also determined that unloading of stalled replisomes requires the activity of p97 segregase, as it is inhibited with p97 inhibitor NMS873 (Fig 5B). From these observations we can thus say that neither prior modification of Mcm7 in S-phase nor the “terminated” conformation of the helicase are essential for mitotic modification of Mcm7 and replisome disassembly.

**Figure 5.**
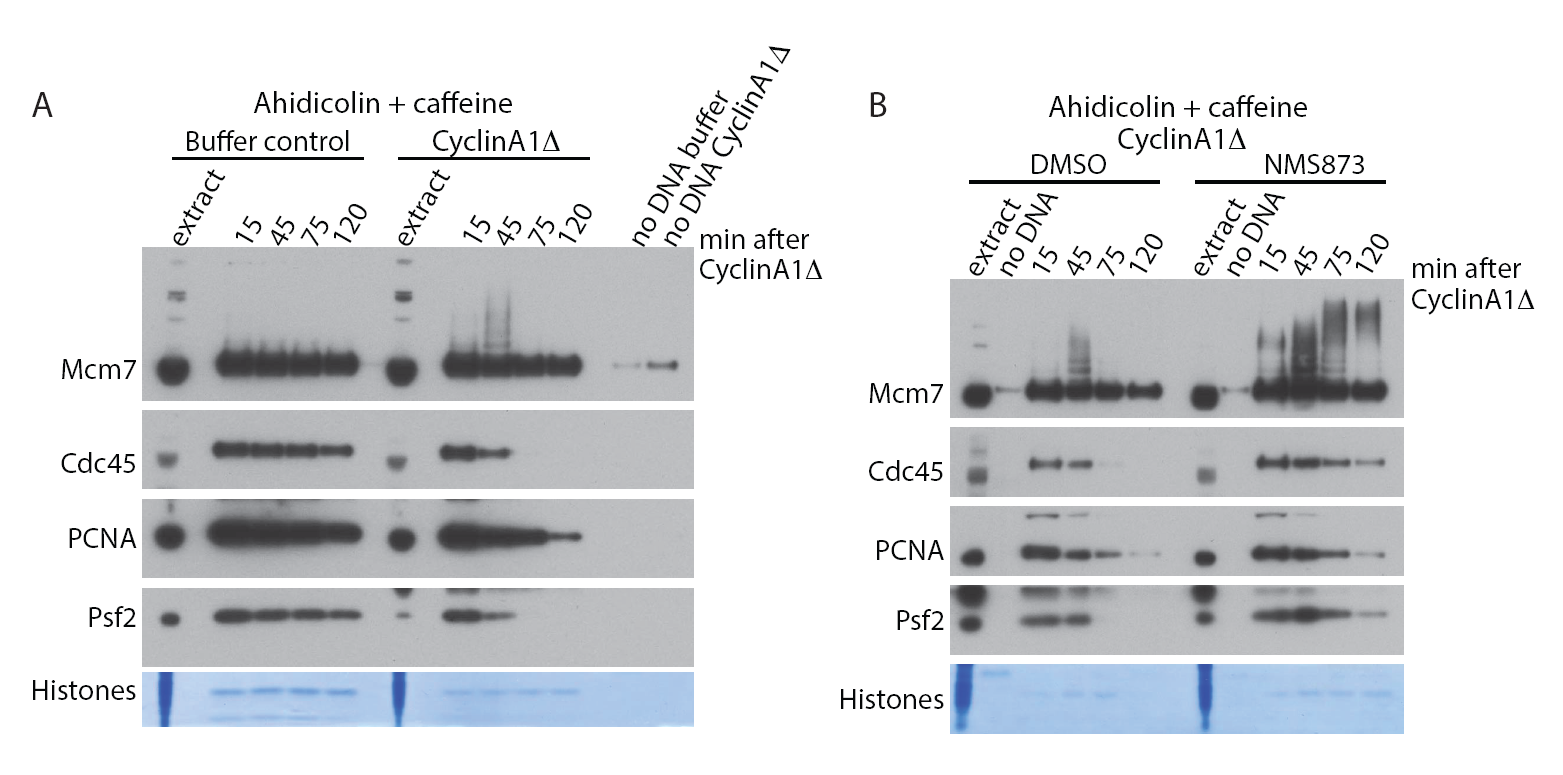
Mitotic unloading of active helicases. **(A)** The replication reaction was performed in egg extract supplemented with DNA polymerase inhibitor aphidicolin and checkpoint inhibitor caffeine. After 90 min of reaction, Cyclin A1Δ was optionally added and chromatin samples isolated at indicated time-points and analysed by western blotting with indicated antibodies (analogously to Fig 1B). **(B)** The inhibition of stalled replisomes was achieved as in (A) and extract was driven into mitosis with optional supplementation with p97 inhibitor NMS873. Chromatin samples were analysed as in (A).

## DISCUSSION

We have presented here the existence of a mitotic pathway of replisome disassembly in vertebrates. One immediate question is why would the cells need a mitotic pathway of replisome disassembly? Traditionally, it is perceived that all the DNA metabolism should be finished before cells enter mitosis. According to this model, G2 phase of the cell cycle is there to ensure that all DNA replication and damage repair are completed prior to chromosome condensation and separation during mitosis. The last decade provided, however, much evidence that this is not the case: unreplicated DNA is detected in many human cells in mitosis; DNA synthesis can proceed during mitosis (mitotic DNA synthesis – MiDAS); underreplicated DNA can lead to the formation of ultrafine bridges (UFB) in anaphase and, finally, formation of structures in G1 stage of the next cell cycle that are bound by 53BP1 protein (53BP1 bodies) (Liu, Nielsen et al., 2014, Minocherhomji, Ying et al., 2015, Moreno, Carrington et al., 2016). Genomewide, such unreplicated regions correlate with common fragile sites (CFS), which are chromosomal loci reponsible for the majority of the rearrangements found in cancer cells (Bhowmick & Hickson, 2017). These unreplicated fragments of DNA result from replication forks not finishing replication and such forks, with their associated replisomes, are subsequently retained on chromatin into mitosis. It is likely that these unreplicated DNA fragments must be processed in mitosis to ensure correct chromosome segregation and this processing will involve replisome unloading and fork remodelling – hence the need for a process of replisome disassembly in mitosis.

TRAIP is a pleiotropic ubiquitin ligase involved in numerous cellular processes. It is clear that TRAIP is essential for appropriate repair of DNA damage in many forms: mitomycin C (MMC) induced inter-strand crosslinks (ICLs); damage caused by treatments with campthotecin (CPT) (Hoffmann et al., 2016), UV (Harley et al., 2016) and hydroxyurea (HU) (Feng et al., 2016); as well as for translesion DNA synthesis (Wallace, Merkle et al., 2014). TRAIP has also been reported to be an important regulator of the spindle assembly checkpoint and regulates mitotic progression. Cells with downregulated TRAIP go through mitosis faster and with more chromosome segregation errors (Chapard et al., 2014, Park, Jo et al., 2015). For most of these processes the ubiquitin ligase activity of TRAIP is essential, but the substrate(s) modified by TRAIP is not known.

In support of our observation that TRAIP interacts weakly with S-phase chromatin when replication forks replicate DNA (Fig 3A), TRAIP has been shown to interact with nascent DNA in unperturbed S-phase in human cells through Nascent Chromatin Capture (NCC) (Hoffmann et al., 2016) but TRAIP knockdown does not affect much replication progression and overall DNA synthesis rates (Harley et al., 2016, Hoffmann et al., 2016). Upon DNA damage TRAIP re-localises from nucleoli to sites of damage in a manner dependent on a PCNA interacting box (PIP-box), present at the C-terminus of TRAIP (Feng et al., 2016, Hoffmann et al., 2016). Loss of TRAIP was suggested to interfere with the reconfiguration of stalled replication forks (Hoffmann et al., 2016), possibly through unloading of PCNA as further inhibition of proteasomal degradation in the absence of TRAIP did not exacerbate the levels of HU-induced fork stalling. This suggested that degradation of a TRAIP ubiquitylation substrate is not the cause of this phenotype (Feng et al., 2016, Hoffmann et al., 2016). Interestingly, cells expressing the ΔRING mutant of TRAIP as the only TRAIP version, are as sensitive to MMC as TRAIP knockdown cells, while ΔPIP TRAIP cells are only mildly sensitive. This indicates that even without PCNA interaction TRAIP can still find its targets at the replication forks (Hoffmann et al., 2016). With the data presented here, identifying TRAIP as the ubiquitin ligase needed for Mcm7 ubiquitylation during mitosis, it is interesting to speculate that TRAIP can play an analogous role during DNA damage repair i.e. to stimulate replisome unloading and fork remodeling.

If TRAIP interacts with replisome and post-termination replisome during normal S-phase (data presented here and (Hoffmann et al., 2016)) an important question emerges, which is how is it activated only in mitosis or only upon DNA damage? In human cells, the abundance of TRAIP protein is tightly regulated and peaks in G2/M stages of cell cycle (Chapard et al., 2015). Moreover, TRAIP re-localizes from nucleoli to sites of DNA damage upon treatment, but we do not know what is driving this change in location. In early mitosis TRAIP is localized all over nucleus but binds exclusively to chromosomes in anaphase (Chapard et al., 2014). The abundance, localization and activity of TRAIP are therefore very carefully regulated in the cell. The intricacy and importance of this regulation may stem from the fact that uncontrolled unloading of the replisome would have a big impact on cell viability and is supported by observations whereby even moderate levels of TRAIP overexpression were reported to be cytotoxic, while epitope tagging of TRAIP (especially at the N-terminus) affects its functionality (Hoffmann et al., 2016).

Our data is consistent with a model in which TRAIP drives mitotic replisome disassembly by promoting Mcm7 modification with K6 and K63 linked ubiquitin chains. Although there is no previous experimental evidence that TRAIP can support such ubiquitin linkages *in vivo, in vitro* assays have shown that TRAIP works well with conjugating enzymes (E2s) UbcH5a,b,c (but not UbcH2, H3, H6, H7, or Ubc13+Uev1A) (Besse et al., 2007). Interestingly, UbcH5a was shown to support formation of ubiquitin chains with no specific topology (Windheim, Peggie et al., 2008). It is plausible therefore that TRAIP/UbcH5 can effectively produce chains of different linkages to support mitotic replisome disassembly.

Upon Mcm7 ubiquitylation the replisome needs to be unloaded. In S-phase, K48 ubiquitin chains support replisome disassembly. In mitosis however, K48 chains are not functional and unloading is driven instead by K6 and K63 chains. We know that p97, in complex with Ufd1 and Npl4 cofactors, is responsible for unloading of the replisome in S-phase (Maric, Mukherjee et al., 2017, Moreno et al., 2014, Sonneville et al., 2017). While p97 is well known for processing substrates ubiquitylated with K48 linked ubiquitin chains (Meyer, Bug et al., 2012), less is known about its contribution in processing other ubiquitin linkages. Interestingly, a recent study shows that upon inhibition of p97 activity, human cells accumulate K6, K11, K48 and, to a lesser extent, K63 ubiquitin chains (Heidelberger, Voigt et al., 2018). Moreover, out of five tested p97 cofactors, all were found to associate with K11 chains, four with K48 chains, and three with K63 chains (Alexandru, Graumann et al., 2008). p97 cofactors are known also to interact with ubiquitin-like modifiers e.g. Nedd8 and Atg8 (reviewed in (Meyer, 2012)). Finally, p97 was also shown to bind more readily to branched K11-K48 chains, than to K11 or K48 chains on their own (Meyer & Rape, 2014). These proteome-wide data imply that the role of p97 does indeed extend beyond recognition of K48-chain-modified substrates, though currently little is known about its interaction with K6 chains.

Importantly, other p97 substrates have been identified, which are modified with non-K48 chains e.g. in double stand break repair RNF8 multi-monoubiquitylates L3MBTL1, which is then extracted by p97 (Kato, Nakajima et al., 2014); in mitochondrial fusion, Mfn1 is ubiquitylated with K63 chains and processed by p97 (Mukherjee & Chakrabarti, 2016) and in antiviral signaling the interaction between p97 and the RIG-I sensor of viral DNA is stimulated by K63 ubiquitylation of RIG-I (Hao, Jiao et al., 2015). All in all, there is much more complexity to p97 function regulation than generally appreciated and future research will reveal how the fine tuning of replisome disassembly by p97 at different stages of the cell cycle or from different DNA structures is achieved.

Finally, we have shown that in the *Xenopus* system neither S-phase nor mitotic replisome disassembly requires SUMO modifications (Fig 4 and EV Fig 3) in contrast to *C. elegans* embryos where ULP-4 is required for mitotic unloading (Sonneville et al., 2017). This requirement may be thus specific to worm embryos, require ULP-4 protein but not its enzymatic activity or regulate an indirect process that is not well recapitulated in the egg extract cell free system. It has been suggested recently that SUMOylation of TRAIP can regulate its stability and ability to move to the nucleus (Park, Han et al., 2016), but this may not be present in the egg extract.

Perturbations in DNA replication initiation and elongation leading to genomic instability are well linked with genetic disorders and can drive cancer development. The disruption of replisome disassembly is, therefore, highly likely to be detrimental to human health too. While so far we have no solid data to support this claim, previous studies with TRAIP do suggest this to be the case: homozygous TRAIP knockdown mouse embryos die shortly after implantation due to proliferation defects (Park et al., 2007); mutations in human TRAIP lead to primordial dwarfism (Harley et al., 2016); overexpression of human TRAIP has been reported in basal cell carcinomas (Almeida, Ryser et al., 2011) and breast cancer (Yang, Trent et al., 2006, Zhou & Geahlen, 2009) and reduced nuclear expression of TRAIP was associated with human lung adenocarcinoma (Soo Lee, Jin Chung et al., 2016). The fact that cells have evolved multiple pathways to ensure timely replisome disassembly supports the notion of the vital importance of this process for cell biology and time will tell whether targeting Mcm7 and replisome disassembly in mitosis is the key mechanism leading to any of these disease phenotypes.

## MATERIALS AND METHODS

### Inhibitors

Caffeine (C8960, Sigma) was dissolved in water at 100 mM and added to the extract along with demembranated sperm nuclei at 5 mM. MLN4924 (A01139, Active Biochem) was dissolved in DMSO at 20 mM and added to the extract 15 minutes after addition of sperm nuclei at 10 μM. NMS873 (17674, Cayman Chemical Company) was dissolved in DMSO at 10 mM and added to the extract 15 minutes after addition of sperm nuclei at 50 μM. SUMO2-VS (UL-759) was purchased from Boston Biochem and used at 1 μM in *Xenopus laevis* egg extract. Aphidicolin was dissolved in DMSO at 8 mM and added to the extract along with demembranated sperm nuclei at 40 μM.

### Recombinant proteins

Recombinant His-tagged ubiquitin and ubiquitin mutants were purchased from Boston Biochem, dissolved in LFB1/50 (40 mM Hepes/KOH pH 8.0, 20 mM potassium phosphate pH 8.0, 50 mM potassium chloride, 2 mM magnesium chloride, 1 mM EGTA; 10% sucrose w/v; 2 mM DTT; 1 μg/ml aprotinin; 1 μg/ml leupeptin; 1 μg/ml pepstatin) buffer at 10 mg/ml and used at 0.5 mg/ml in *Xenopus laevis* egg extract. pET28a-X.l. SUMO1 and pET28a-X.l. SUMO2 were purchased from GenScript. Recombinant His-tagged *Xenopus laevis* SUMO1 and SUMO2 were expressed in Rosetta (DE3) pLysS cells over night at 20°C after induction with 1 mM IPTG. Cells were lysed in lysis buffer: 50 mM Tris-HCl, 500 mM NaCl, 10 mM imidazole, 2 mM MgCl_2_, 0.1 mM PMSF, 1μg/ml of each aprotinin, leupeptin and pepstatin, pH 7.5. Homogenates were supplemented with 25 U/ml benzonase and incubated at room temperature for 20 minutes. Homogenates were subsequently spun down at 14,000 g for 30 minutes at 4°C and supernatants incubated with 2 ml of pre-washed Super Ni-NTA Affinity Resin (SUPER-NINA100, Generon) for 2 hours with rotation at 4 °C. Resins were subsequently washed twice with 50 mM Tris-HCl, 500 mM NaCl, 30 mM imidazole, 0.1 mM PMSF, 1μg/ml of each aprotinin, leupeptin and pepstatin, pH 7.5. Resin-bound proteins were finally eluted in 1 ml fractions with a solution containing 50 mM Tris-HCl, 150 mM NaCl, 200 mM imidazole, 5 mM β-mercaptoethanol, 0.1 mM PMSF, 1 μg/ml of each aprotinin, leupeptin and pepstatin, pH 7.5. Fractions containing the highest levels of recombinant SUMO1 or SUMO2 were dialysed into LFB1/50 buffer. Both SUMO1 and SUMO2 were used at 0.5 mg/ml in *Xenopus laevis* egg extract.

pET28a-pHISTEV30a-SENP1(415-649) was a kind gift from Ron Hay’s laboratory. Recombinant active domain of human SENP1 (aa 415-647) was expressed and purified as explained above for recombinant SUMOs.

Recombinant His-tagged *Xenopus laevis* cyclin A1 NΔ56 (pET23a-*X.l.* cyclin A1 NΔ56) was a kind gift from Julian Blow’s laboratory (Strausfeld et al., 1996), was expressed in Rosetta (DE3) pLysS cells over night at 15°C after induction with 1 mM IPTG, and subsequently purified as explained above for recombinant SUMOs but using different solutions. Lysis buffer: 50 mM Tris-HCl, 300 mM NaCl, 2 mM MgCl_2_, 1 mM DTT, 0.1 mM PMSF, 1 μg/ml of each aprotinin, leupeptin and pepstatin, pH 7.4. Washes: Resin was washed twice with lysis buffer on its own and twice again with lysis buffer supplemented with 0.1% Triton X-100.

Elution buffer: Lysis buffer supplemented with 10% glycerol and 250 mM imidazole. *Xenopus* TRAIP was cloned into pGS21 vector, expressed in BL21 (DE3) bacterial strain in autoinduced AIM media O/N at 18°C. Pellets were lysed in lysis buffer: 50mM NaH2PO4; pH 9, 300mM NaCl, 10% glycerol, 2mM DTT, 2mM MgCl_2_, 0.05% Brij, 0.1 mM PMSF, 1 μg/ml of each aprotinin, leupeptin and pepstatin, 1 mg/ml lysozyme, 25 U/ml Benzonase. The protein was purified as above but using Glutathione Sepharose 4B (GE Healthcare) and eluted with 25 mM glutathione. The protein was then dialised into LFB1/50 buffer (as above) and concentrated up to 0.3 mg/ml of full length GST-TRAIP. It was used in the egg extract at final concentration of 30 μg/ml. pGS21-TRAIP(C25A) was generated by site directed mutagenesis and purified in analogous way.

### Antibodies

α-PCNA (P8825) and α-HIS (H1029) were purchased from Sigma. α-TRAIP (NBP1-87125) and α-RNF213 (NBP1-88466) were purchased from Novus Biologicals. α-SUMO2 and α-SUMO1 were produced in the lab by culturing the hybridoma cell line SUMO2 (8A2) and SUMO1 (21C7) purchased from Developmental Studies Hybridoma Bank (hybridoma cell culture was done following manufacturer’s instructions and adding 20 mM L-glutamine to the media). Affinity purified α-Cdc45 (Gambus, Khoudoli et al., 2011), α-Mcm3 (Khoudoli, Gillespie et al., 2008) and α-LRR1 (S962D) (Sonneville et al., 2017) were previously described. α-Mcm7 was raised in sheep against recombinant *Xenopus laevis* Mcm7, purified from *E.coli* and affinity purified in the lab.

### DNA staining and microscopy

Interphase *Xenopus laevis* egg extract was supplemented with 10 ng/μl of demembranated sperm nuclei and incubated at 23°C for 90 minutes to allow completion of DNA replication. Mitosis was optionally driven by addition of 826 nM cyclin A1 NΔ56, and reactions were incubated for further 2 hours. Assembled S-phase or mitotic chromatin were stained with Hoechst 33258 and viewed as previously described (Strausfeld et al., 1996).

### DNA synthesis assay (TCA)

The replication reactions were started with the addition of demembranated *Xenopus* sperm DNA to 10 ng/µl as described before (Gillespie et al., 2012). The synthesis of nascent DNA was measured by quantification of α32P-dATP incorporation into newly synthetized DNA as described before (Gillespie et al., 2012).

### Chromatin isolation time-course

Interphase *Xenopus laevis* egg extract was supplemented with 10 ng/μl of demembranated sperm DNA and subjected to indicated treatments. The reaction was incubated at 23°C for 90 minutes to allow completion of DNA replication, after which mitosis was driven by addition of 826 nM cyclin A1 NΔ56. The extract was optionally supplemented with inhibitors or recombinant proteins along with cyclin A1 NΔ56 as indicated. Mitotic chromatin was isolated in ANIB100 buffer supplemented with 10 mM 2-chloroacetamide (Millipore) and 5 mM N-ethlylmaleimide (NEM) (Acros Organics) at indicated time-points after addition of cyclin A1 NΔ56 as previously described (Gillespie et al., 2012).

### Immunoprecipitation of post-termination CMG associated with mitotic chromatin

3.75 ml of interphase *Xenopus laevis* egg extract was supplemented with 10 ng/μl of demembranated sperm nuclei, 5 mM caffeine and 50 μM p97 inhibitor NMS873. The reaction was incubated at 23°C for 90 minutes to allow completion of DNA replication, after which mitosis was driven by addition of recombinant cyclin A1 NΔ56 at 826 nM followed by incubation at 23°C for further 60 minutes. At this stage, chromatin was isolated in ANIB100 buffer supplemented with 10 mM 2-chloroacetamide as previously described (Gambus et al., 2011) and chromatin bound protein complexes released into solution by chromosomal DNA digestion with 2 U/μl benzonase for 15 minutes. Solubilisation of chromatin bound protein complexes was further facilitated by subjecting the sample to 5 minutes of 30 seconds ON/OFF sonication cycles using a diagenode bioruptor and increasing the concentration of potassium acetate up to 150 mM. The resulting protein complexes were subsequently subjected to either non-specific IgG (from sheep serum) or Mcm3 immunoprecipitation and the immunoprecipitated material analysed by mass spectrometry as previously described (Sonneville et al., 2017) in collaboration with Dr Richard Jones from MS Bioworks LLC.

## ACKNOWLEDGMENTS

Dr Sara Priego Moreno was funded by Wellcome Trust ISS Award, Dr Rebecca Jones and Dr Agnieszka Gambus were funded by MRC CDA MR/K007106/1. We would like to thank Prof Ron Hay for pET28a-pHISTEV30a-SENP1(415-649) and Prof Julian Blow for pET23a-*X.l.* cyclin A1 NΔ56.

---

**EV Figure 1.**
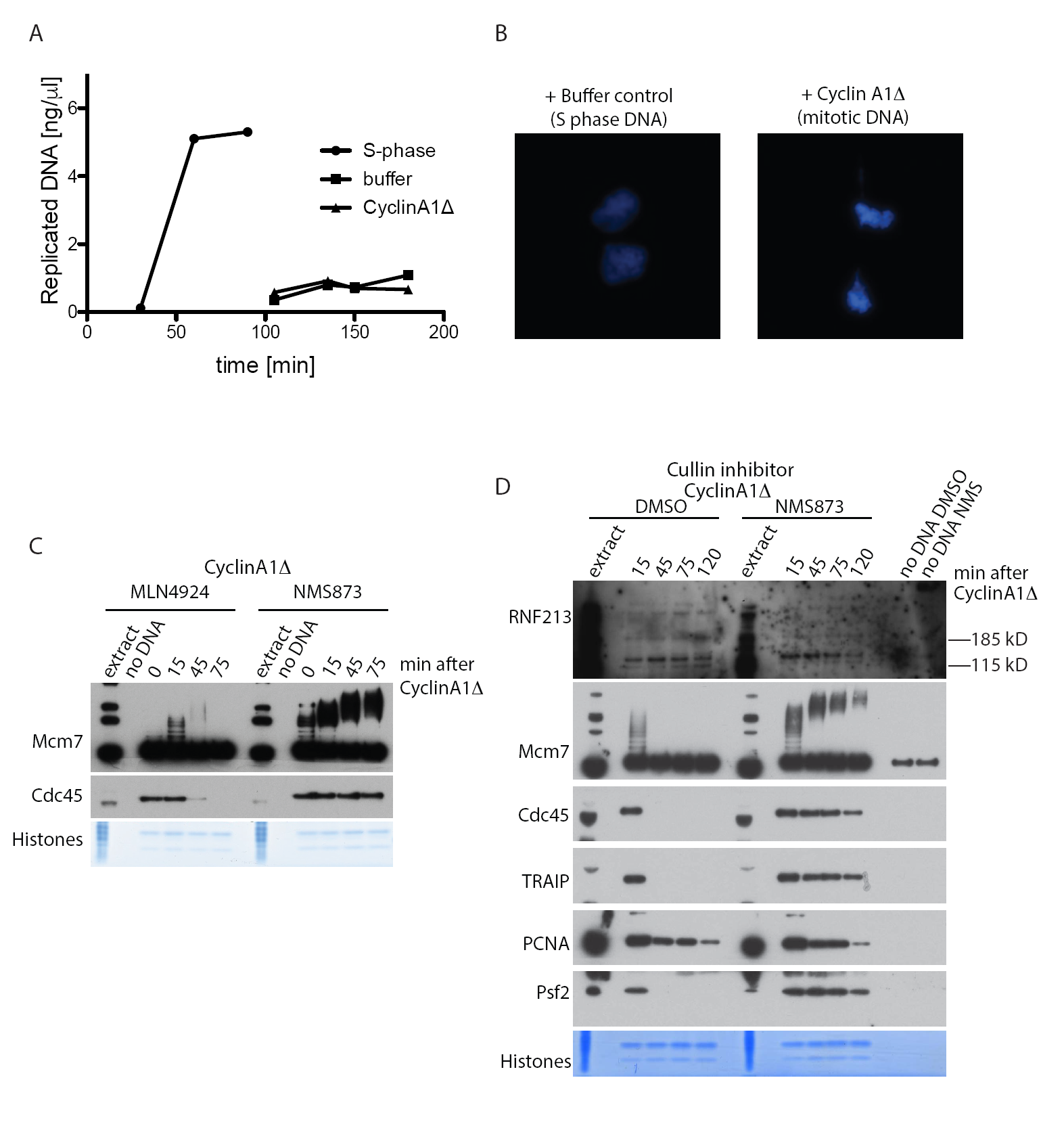
Setting up mitotic replisome disassembly. **(A)** Progression of replication reaction was measured by incorporation of radioactive dATP into the DNA. P^32^α-dATP was added to the extract at the beginning of the reaction (S-phase) or after 90 min of replication reaction together with optional addition of Cyclin A1Δ (buffer control or Cyclin A1Δ). **(B)** The replication reaction was completed and optionally driven into mitosis by addition of Cyclin A1Δ. The nuclei assembled in the extract were visualised by Hoechst 33258 staining. **(C)** Experiment as in Fig 1C but indicated inhibitors were present in the reaction throughout both stages of the cell cycle.

**EV Figure 2.**
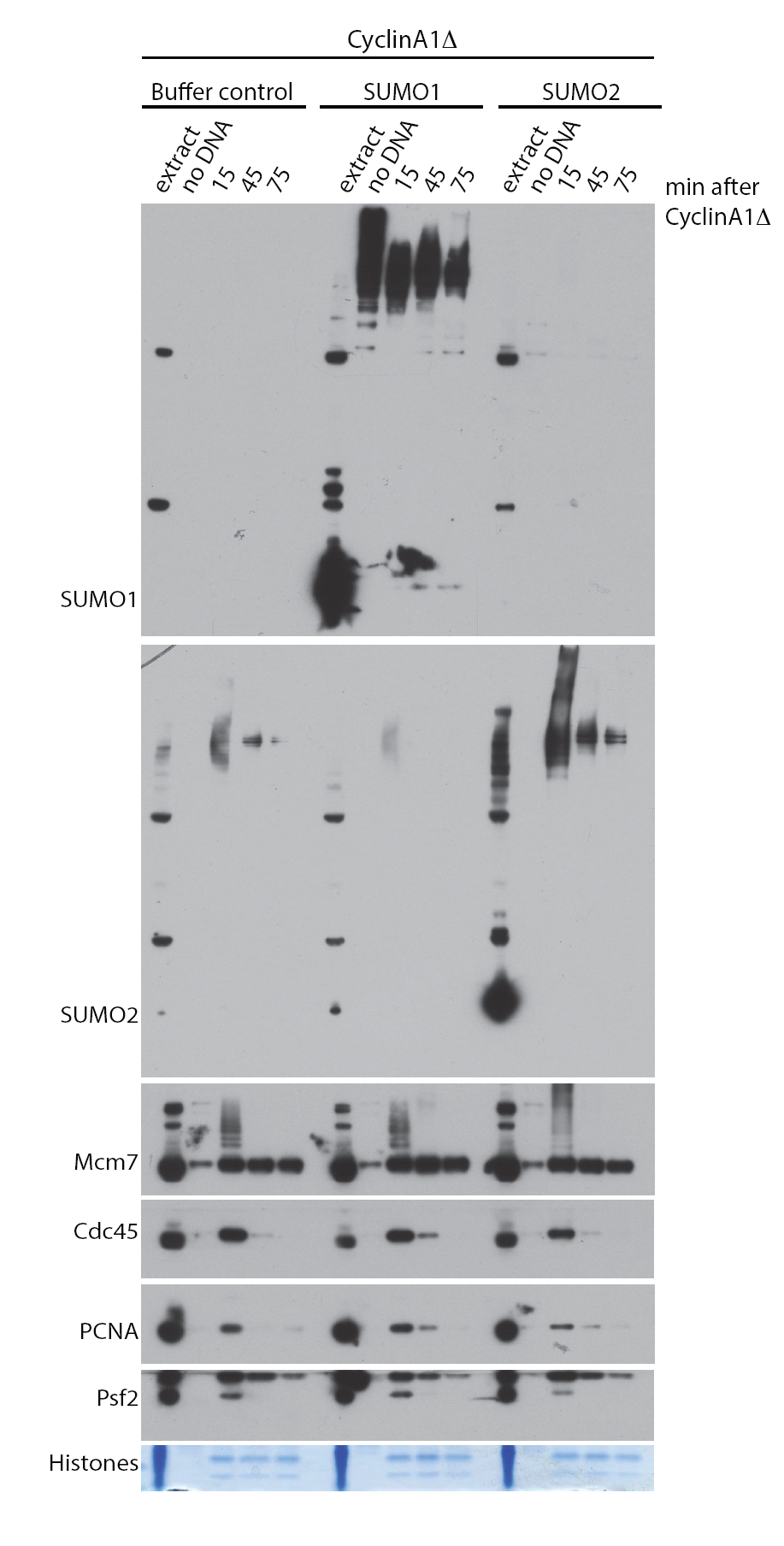
Mitotic replisome disassembly is not affected by stimulation of SUMOylation. The replication reaction was completed in the presence of Cullin inhibitor MLN4924 and driven into mitosis by addition of Cyclin A1Δ. At the same time as Cyclin A1Δ, extract was optionally supplemented with buffer control, SUMO1 or SUMO2. Chromatin samples were isolated at indicated time-points and analysed as in Fig 1.

**EV Figure 3.**
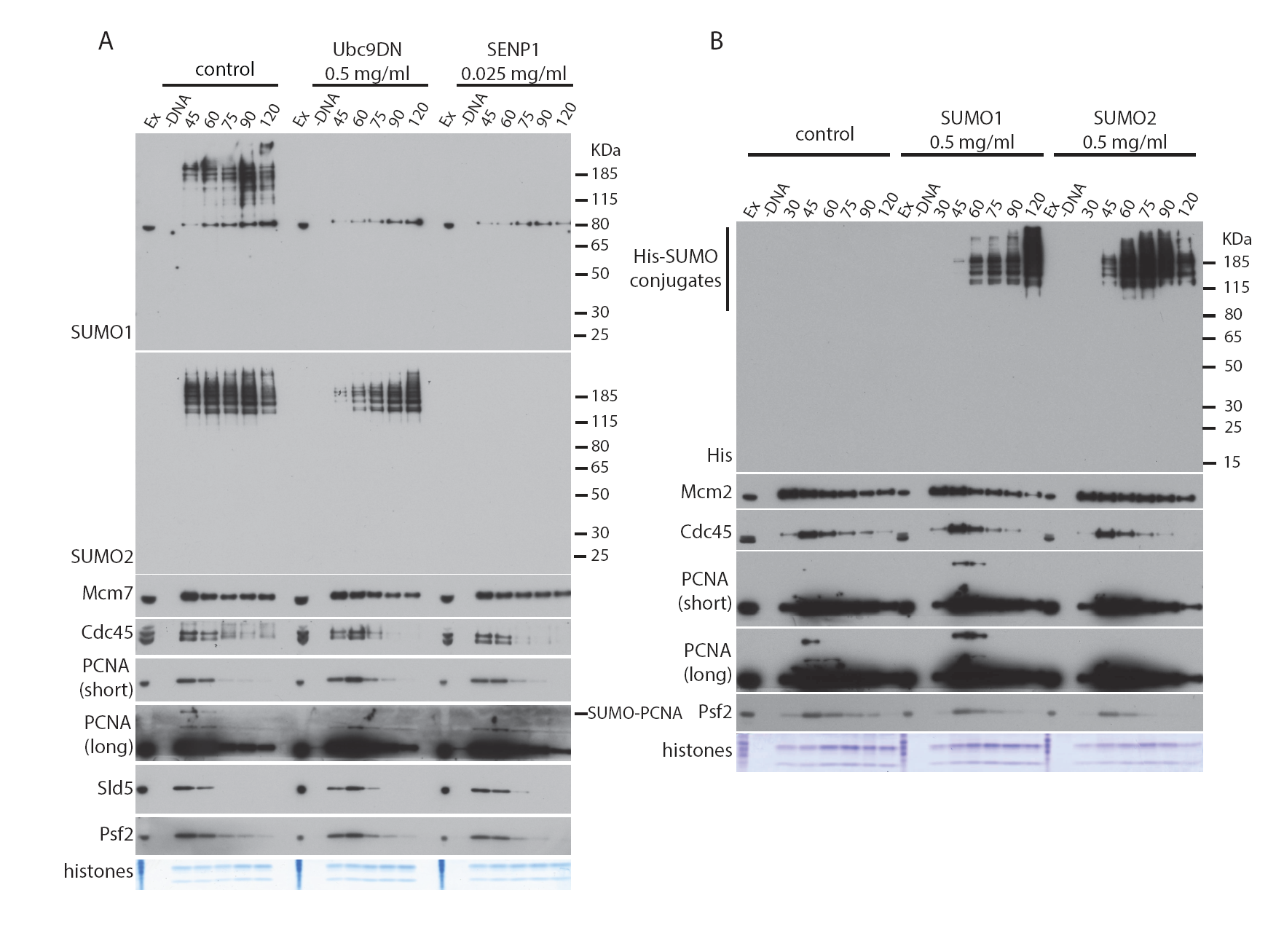
S-phase replisome disassembly is not affected by SUMOylation. **(A)** The replication reaction was performed in egg extract supplemented with dominant negative mutant of Ubc9 (Ubc9DN) or active domain of SENP1. Chromatin samples were isolated at indicated time-points and analysed by western blotting with indicated antibodies. **(B)** The replication reaction was performed in egg extract supplemented with 0.5 mg/ml of SUMO1 or SUMO2 (as indicated). Chromatin samples were isolated at indicated time-points and analysed by western blotting with indicated antibodies. Controls as in Fig 1.

